# Using iPALM to determine protein organisation in cardiac muscle Z-discs

**DOI:** 10.64898/2026.05.08.723761

**Authors:** Oliver Umney, Alistair P. Curd, Heather L. Martin, Tarek Lewis, Anna Tang, Harikrushnan Balasubramanian, Satya Khuon, Jesse Aaron, Michelle Peckham

**Affiliations:** School of Molecular and Cellular Biology, Faculty of Biological Sciences, University of Leeds, Leeds, LS2 9JT UK; Engineering and Physical Sciences Faculty Services, Faculty of Engineering and Physical Sciences, University of Leeds, Leeds LS2 9JT UK; Advanced Imaging Center, Janelia Research Campus, Howard Hughes Medical Institute, Ashburn, VA 20147, USA; Astbury Centre for Structural and Molecular Biology University of Leeds, Leeds LS2 9JT UK

**Keywords:** iPALM, super-resolution microscopy, Adhiron, Z-disc, Striated muscle

## Abstract

Sarcomeres, the basic repeating unit of striated muscle, are joined together by crosslinked actin filaments found at the boundaries of muscle sarcomeres, termed Z-discs. Z-discs play a key role in cardiac signalling and disease, however, the arrangement and function of many of the proteins present in the Z-disc remain to be understood. Here, we determined the organisation of 3 key proteins, ZASP, ɑ-Actinin-2 and the Z1Z2 epitope of titin, located within the Z-disc. We fluorescently labelled these proteins in cardiac myofibrils using Adhirons specific to each protein and used interferometric photoactivated localization microscopy (iPALM) to obtain the 3D position of these proteins to a high precision (<10nm in x,y,z). We then used PERPL (Pattern Extraction from Relative Positions of Localisations) to analyse patterns in the relative positions of the proteins and reveal their underlying organisation. This analysis revealed that ZASP and ɑ-Actinin-2 have a similar repeating organisation, but that the organisation of Z1Z2 is different.

## Introduction

Sarcomeres are the basic repeating unit of striated muscle. In this type of muscle, sarcomeres are linked end-to-end by structures known as Z-discs, to generate roughly cylindrical myofibrils that run from one end of the cell to the other. Precision building of the constituent proteins means that each muscle sarcomere is highly similar to every other sarcomere in the muscle cell. This is important in ensuring each sarcomere generates equivalent force and movement.

Proteins from over 100 genes have been linked to the Z-disc [1], of which more than half localise to its core. These proteins can have either structural or signalling roles, or both. Of these proteins, ɑ-actinin plays a key role in crosslinking and anchoring actin-filaments from adjacent sarcomeres within the Z-disc (Luther, 2009). Others, such as filamin C, myotilin and ZASP, play both structural and signalling roles through their interaction with other proteins (reviewed in [2, 3]). Z-disc sensing of mechanical strain and subsequent downstream signalling can remodel the Z-disc and muscle sarcomeres and is important in both vertebrate and invertebrate [4] Understanding how Z-disc proteins are organised and interact with each other is crucial not only to understanding their biological function, but in understanding disease processes. Despite this, we still know very little about how proteins are organised within the Z-disc.

One of the main challenges in uncovering protein organisation in the Z-disc is that it is both narrow and variable in structure. At its widest in cardiac muscle, it is still only approximately 100 to 140 nm [5]. The lattice structure within the Z-disc has even smaller dimensions of approximately 25 nm in x, y and z [5]. These dimensions are all below the resolution limit of the light microscope (250 nm). Moreover, the lattice structure can either take a small-square (predominantly relaxed muscle) or basketweave form (predominantly contracting muscle) with some mixture of the two typically present [5–7]. The transition from small square to basketweave has been suggested to involve a change in twist in ACTN2, accompanied by an increase of ∼ 10-20% in lattice dimensions. This is likely because the interaction between the calponin homology (actin-binding) domain of ACTN2 and F-actin is highly flexible [7–9]. However, in rigor muscle, the lattice always takes the basketweave form [10]. Even the improved resolution of electron microscopy has failed to reveal the organisation of Z-disc proteins. Sub-tomogram averaging blurs out the detail in the Z-disc structure, and specific proteins cannot be identified (excepting actin and α-actinin) [11–13]. Therefore, other approaches to uncovering its nanoscale organisation need to be tested and developed.

Super-resolution imaging, using single molecule localisation microscopy approaches, has proved to be a powerful approach to determine the organisation of multiple proteins within focal adhesions [14], endocytic vesicles [15] and other dense cytoskeletal structures (reviewed in [16]). These approaches generally obtain resolutions of approximately 20 nm in 2D (x,y), but resolution in the third dimension (z) is at best approximately 50 nm (reviewed in [16]). Interferometric photoactivated light microscopy (iPALM), which combines single molecule localisation with multiphase interferometry, achieves a much higher resolution in this dimension of approximately 10 nm [17]. Thus, the resolving power of iPALM is ideally suited to investigate the organisation of proteins within the Z-disc, as it closely matches the dimensions of the Z-disc lattice.

To use iPALM to image protein organisation within the Z-disc, typically antibodies would be used to label specific proteins. However, the combination of both primary and secondary antibodies commonly used to do this poses two problems. First, the dye, which is attached to the secondary antibody, can be over 20 nm from the target site, as both antibodies are approximately 10 nm in size. This is compounded by the flexibility of the antibodies, which can further blur the dye position [18]. Second, the large size of the antibodies can also restrict their ability to fully label dense cytoskeletal structures, as they can be excluded from their core [19, 20]. To overcome this, we isolated non-antibody binding proteins (Adhirons [21] also known as Affimers) to specific proteins within the Z-disc. These proteins are about 4 nm in size, approximately 10-12 kDa in mass, and contain two binding loops as part of a scaffold, which specifically bind the protein of interest (reviewed in [19, 20]).

Here we have used Adhirons to label cardiac myofibrils, iPALM to image them, and have used downstream software (PERPL [22]) to determine the pattern of organisation of three Z-disc proteins. The Adhirons we used either specifically bind the calponin-homology (actin binding) domains of α-actinin-2 (ACTN2) [22, 23], the N-terminal Z1Z2 epitope of the giant protein titin, or the signalling protein ZASP (Z-band alternatively spliced PDZ motif protein (also termed Cypher and LIM domain-binding protein 3, and PDLIM6). We chose ACTN2, as its organisation within the Z-disc is the best characterised and is expected to show a characteristic pattern. Titin is a giant molecule, up to 4 MDa in size and 1 µm in length, which spans from its N-terminal region within the Z-disc to the C-terminal region in the M-line in the middle of the muscle sarcomere [24]. We chose the Z1Z2 epitope of titin as it has been reported to be located either in the periphery of the Z-disc [25] or in its core [26], and thus the Adhiron could resolve these findings and should reveal its pattern of organisation, if any. We chose ZASP, as its PDZ domain interacts with ACTN2 and immunoEM shows it is distributed throughout the Z-disc [27]. Thus, ZASP might be expected to show a similar pattern of organisation to ACTN2.

## Results

### Adhirons to ACTN2, the Z1Z2 epitope of titin, and ZASP

The Adhirons isolated to the calponin homology (CH) domains of ACTN2, the Z1Z2 domains (N-terminal immunoglobulin tandem domains) of titin and ZASP have been briefly reported on in earlier work from our group [22, 23]. The structure of the co-crystal of the Adhiron bound to the calponin homology (CH) actin binding domains of ACTN2 (6WST) shows its binding is mediated by salt bridges between Lys72 and Asp 80 in loop 1 of the Adhiron, with Glu217 and Lys221, respectively in the second CH domain of ACTN2 [22]. We do not have co-crystal structures for the remaining two Adhirons. However, Alphafold modelling suggests that that the Adhiron to Z1Z2 binds to the Z2 domain and that the Adhiron to ZASP likely binds to the PDZ (postsynaptic density protein (PSD95), Drosophila disc large tumor suppressor (DlgA), and zonula occludens-1 protein (zo-1) domain (Fig. S1). Pull-down assays demonstrated that the Adhirons bound specifically to their target epitopes. As the first few Ig domains of titin within the Z-disc show some sequence conservation, we also confirmed that the Adhiron to Z1Z2 did not bind to downstream Z-repeats (from Ala201 to Tyr598) (Fig S1).

### iPALM generates high-precision localisations for Z-disc proteins in 3D

iPALM was used to image all three Z-disc proteins, ACTN2, Z1Z2 and ZASP in cardiac myofibrils in rigor conditions, using Alexa647-labelled Adhirons (Figure 1). This generated high precision-localisations in 3D for all three proteins, with the greatest precision in z, followed by y then x (Fig. S2). The mean sarcomere length was estimated as 1.8 ± 0.1 µm (mean ± S.D.), in agreement with the expected sarcomere length for cardiac myofibrils.

**Figure 1.**
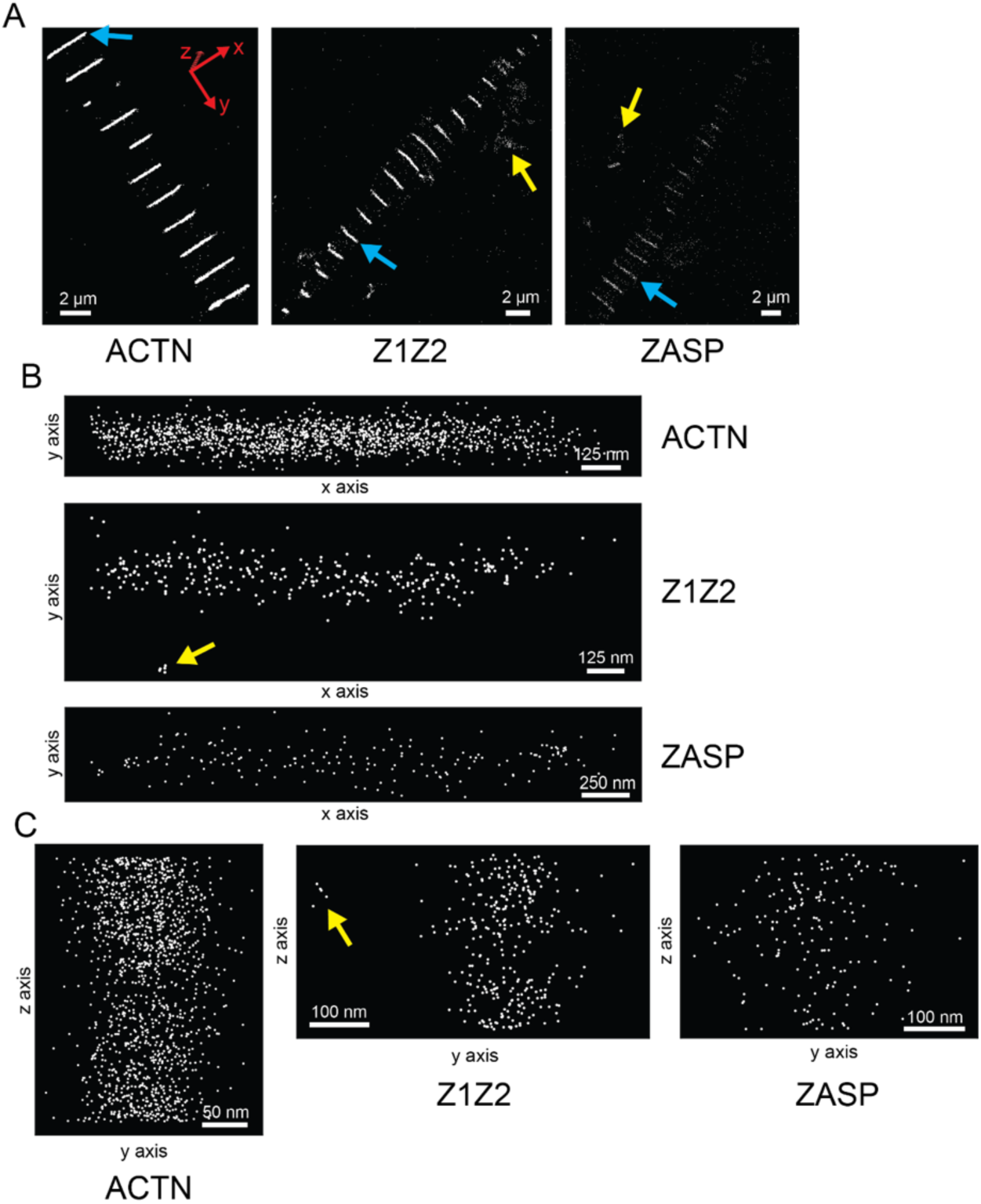
iPALM imaging of Z-disc proteins. **A**: Example iPALM data for ACTN2, Z1Z2 and ZASP in multiple Z-discs along a myofibril. The z axis is into the plane. **B,C**: Localisations for individual Z discs from **A** (blue arrows) in xy (**B**) and yz (**C**) planes. Yellow arrows: localisations removed from further analysis by Z-disc segmentation and denoising. Plots show maximum projection in the orthogonal plane to the view.

Interestingly, the density of localisations for ACTN2 was higher than those for Z1Z2 and lowest for ZASP (Fig. 1A). This may be explained by ACTN2 being the most common Z-disc protein of the three, with ZASP the least.

The localisations for each Z-disc were extracted, allowing for per Z-disc characterisation and for subsequent modelling of protein organisation using PERPL (Figure 1B & C). As the orientation of the myofibrils varies across images, we developed a pipeline to segment the Z-discs and align each one to facilitate downstream pattern analysis (PERPL). Briefly, the iPALM data was converted to images and imported into Ilastik, an interactive semi-automatic segmentation tool, to segment the Z-discs in image space [28]. These segmentations were used to extract the point cloud data for each Z-disc [29], which were then aligned using principal component analysis (PCA) and denoised using DBSCAN (Fig. S3).

As in the unsegmented data, the localisation density for each Z-disc was greatest for ACTN2, followed by Z1Z2 and ZASP (Table 1). The lengths in x and y, together with the measured volume of the Z-discs was largest for ZASP, followed by Z1Z2 and then ACTN2, which may indicate that ZASP has a broader distribution at the Z-disc. Dimensions in z were similar for all three proteins. It could also result from selection bias when segmenting the Z-discs, an artefact of the smaller sample sizes for ZASP and Z1Z2 (Table 1). A further possibility is that the ZASP and Z1Z2 Adhirons were less specific to their target, although data from pulldowns suggest this is not the case (Fig. S1). For ZASP, the particularly small sample size (n_zdiscs_ = 5) resulted from fewer initial iPALM acquisitions and an increased difficulty in segmenting Z-discs from the lower density data.

**Table 1.** iPALM Data characterisation. n_zdiscs_: Number of Z-discs acquired. n_locs_: Number of localisations per Z-disc. L: length of Z-disc in x, y and z. σ: localisation precision in x, y, and z. ρ: localisation density per Z-disc. Values are the mean ± S.D.

| Protein | $n_{\text{locs}}$ | $L_x$<br>($10^3$ nm) | $L_y$<br>($10^2$ nm) | $L_z$<br>($10^2$ nm) | $\sigma_x$<br>(nm) | $\sigma_y$<br>(nm) | $\sigma_z$<br>(nm) | $\rho$<br>($10^3 \mu\text{m}^{-3}$) |
| --- | --- | --- | --- | --- | --- | --- | --- | --- |
| ACTN2<br>($n_{\text{zdiscs}} = 65$) | 1191 ± 762 | 1.84 ± 0.68 | 3.00 ± 0.64 | 2.88 ± 0.09 | 12.7 ± 4.1 | 12.3 ± 4.4 | 10.3 ± 5.2 | 12.7 ± 6.9 |
| Z1Z2<br>($n_{\text{zdiscs}} = 29$) | 353 ± 143 | 2.06 ± 0.53 | 3.35 ± 0.56 | 2.89 ± 0.03 | 13.2 ± 4.5 | 11.9 ± 4.4 | 9.3 ± 4.5 | 3.0 ± 0.8 |
| ZASP<br>($n_{\text{zdiscs}} = 5$) | 134 ± 24 | 2.69 ± 0.23 | 4.55 ± 0.97 | 2.87 ± 0.02 | 11.9 ± 4.7 | 10.1 ± 4.1 | 7.7 ± 3.7 | 0.8 ± 0.2 |

### PERPL reveals the organisation of ACTN2, Z1Z2 and ZASP in Z-disc data

PERPL was used to infer the organisation of ACTN2, Z1Z2 and ZASP in the axial (y) direction and transverse (xz) plane, by modelling the relative position distributions (RPDs) for the localisations (Methods, Fig. S3) [22]. Model RPDs contained different numbers of characteristic distances, which could either be independent of one another, or dependent on one another (such as a linear repeat of a targeted protein domain along one direction). The model RPDs also contain different background models for distances unaccounted for by the combination of characteristic distances (e.g. a spatially random pattern of false positive localisations). For each protein, we swept through and selected pre-processing settings for the localisation data to generate different experimental RPDs. For each RPD, we swept over a range of possible (user-defined models) to fit to the data. We then compared relative likelihoods of the fitted models of axial and transverse organisation (**Tables S1-S2**) [22].

Finally, we selected the fitted model best describing the structure underlying the experimental data in each case, as determined by the AICc (Aikaike criterion) (**Fig. 2**, **Tables 2-3**, **Figures S3-5**). In all cases except for the arrangement of Z1Z2 in the transverse plane, the best model was more than 100 times more likely as an explanation of the data than the second best as determined.

**Figure 2.**
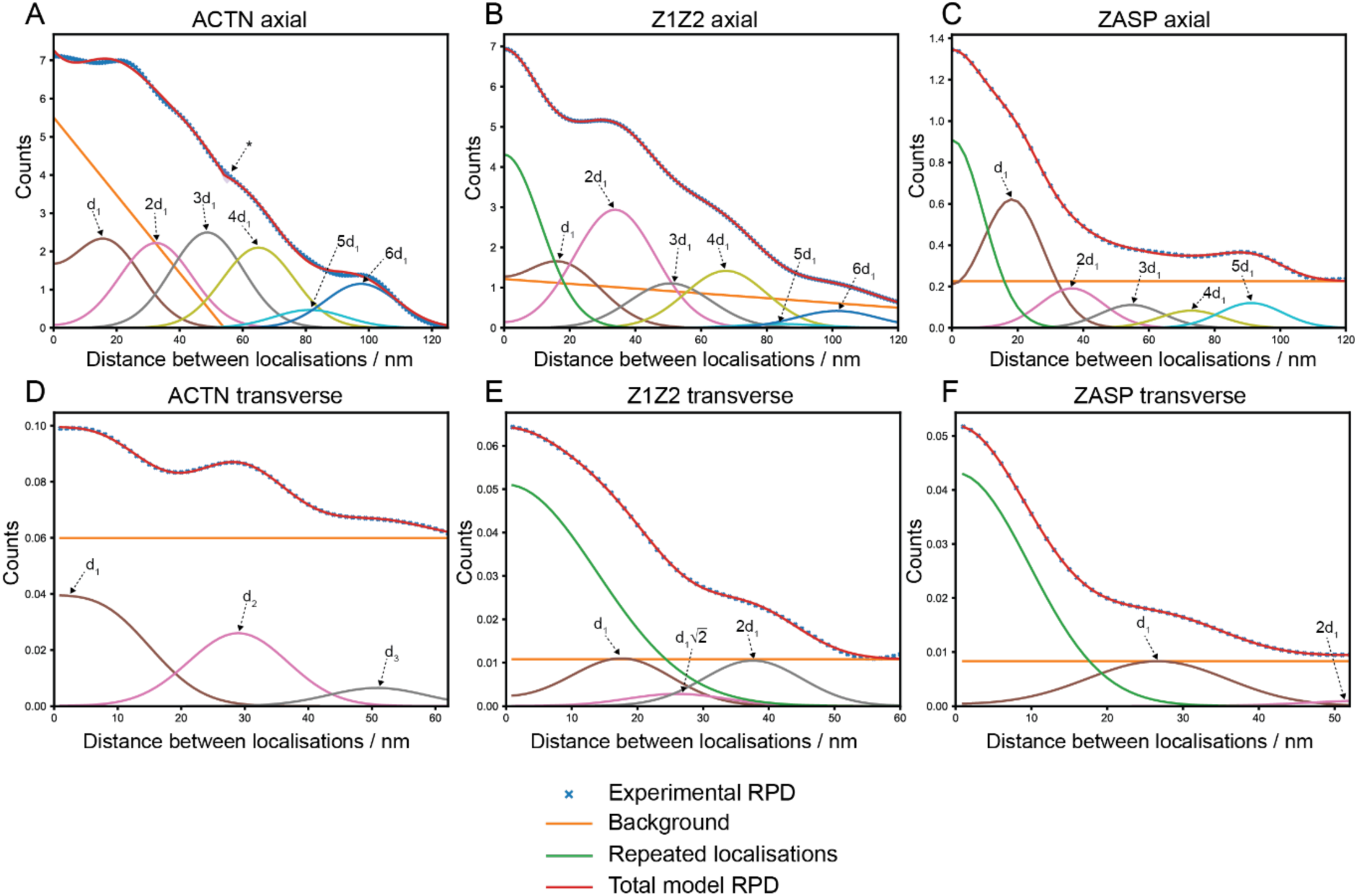
Experimental and best model Relative Position Distributions (RPDs). Total model RPD contains terms for the background from randomly distributed localisations, repeated localisation of the same dye molecule and the characteristic distances (d_1_, d_2_, d_3_) between localisations (**Tables 2,3**). Ratios between modelled distances are shown (e.g. 2d_1_, = 2 * d_1_). 95% confidence intervals shown on total model RPD, only noticeable in A (*).

**Table 2.** Characteristic axial distances and broadening in most likely models. Next most likely model in each case had a relative likelihood of < 0.01.

| Protein | Repeat distance<br>($d_1$, nm) | $\sigma_{\text{broadening}}$ (nm) |
| --- | --- | --- |
| ACTN2 | $16.25 \pm 0.09$ | $11.47 \pm 0.33$ |
| Z1Z2 | $16.88 \pm 0.05$ | $12.40 \pm 0.37$ |
| ZASP | $18.22 \pm 0.06$ | $9.68 \pm 0.11$ |

**Table 3.** Characteristic transverse distances and broadening in most likely models. ͕͐ corresponds to previously reported distance between parallel actin filaments (*d*_p-fil_). ͔͐ corresponds to expected distance between anti-parallel actin filaments (*d*_p-fil_ / √2). ^a^ √2 × d_1_, ^b^ 2 × d_1_.

| Protein | $d_1$ (nm) | $d_2$ (nm) | $d_3$ (nm) | $\sigma_{\text{broadening}}$ (nm) |
| --- | --- | --- | --- | --- |
| ACTN2 | $10.32 \pm 0.03$ | $30.0 \pm 0.06 \zeta$ | $51.41 \pm 0.22$ | $7.65 \pm 0.04$ |
| Z1Z2 | $19.14 \pm 0.23 \zeta$ | $27.07 \pm 0.33^a \zeta$ | $38.28 \pm 0.46^b$ | $7.37 \pm 0.33$ |
| ZASP | $28.17 \pm 0.02 \zeta$ | $56.34 \pm 0.46^b$ | - | $8.96 \pm 0.05$ |

In the axial direction (Fig. 2A-C), we found a linear repeat of between 16 and 18 nm to be the best model for the arrangement of all three proteins, with a variability (σ_broadening_, fitted s.d. of the component peak) of 10–12 nm on the characteristic distances between instances of the same protein (**Table 2**, **Fig. 3A**). This arrangement is expected for ACTN2 from the known structure of the Z-disc complex and its ACTN2 repeated crosslinks [5, 13]. The titin Z1Z2 domains and ZASP therefore apparently follow the same pattern as ACTN2 in the axial direction. However, for ZASP localisations, the signal (peak amplitude) decreased more significantly at higher multiples of the repeat distance. Thus, although we fit 5 peaks, the localisations are more likely to be separated by only one repeat of the Z-disc structure (18 ± 10 nm, mean distance ± σ_broadening_) (**Fig. 2C**, **Table S5**) in agreement with the lower density of localisations (**Table 1**). One possible explanation for this is that the PDZ domains of ZASP, labelled by the Adhiron, tend to be found in pairs (possibly resulting from ZASP dimerization) separated by 16-18 nm, but the occurrence of this pairing occurs randomly throughout the Z-disc.

**Fig. 3.**
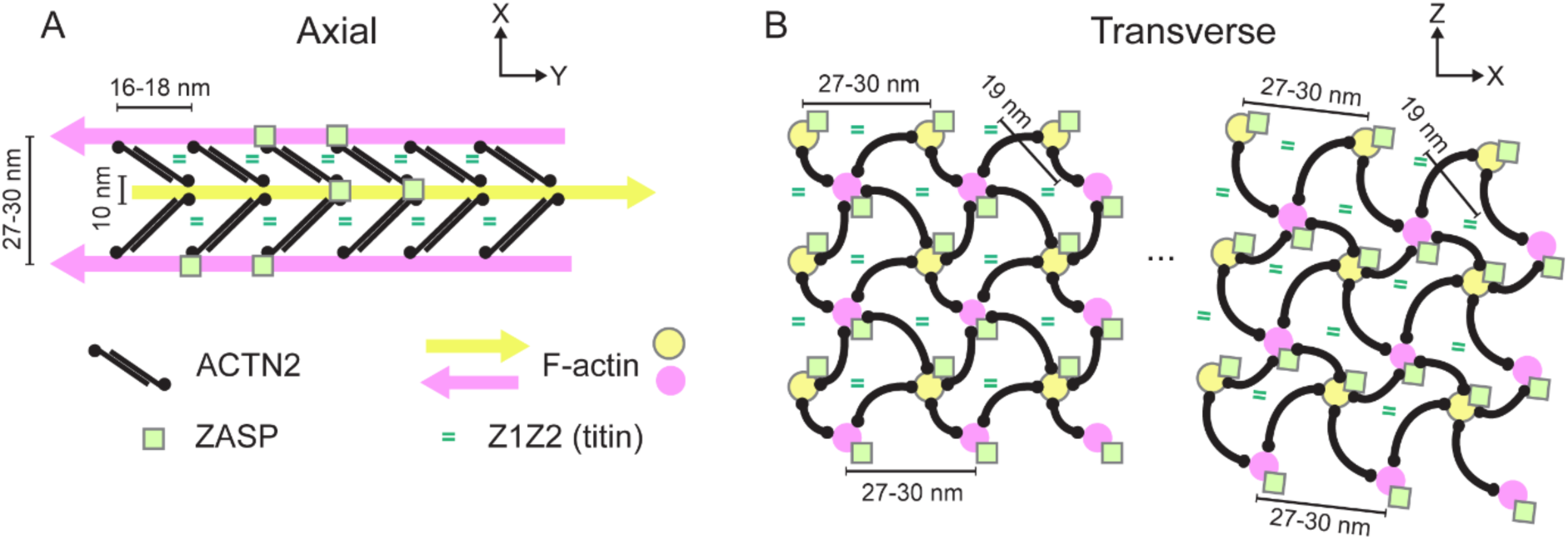
Model for organisation of ACTN2, ZASP and titin Z1Z2 domains in the Z-disc (cardiac myofibrils). **A:** Arrangement of actin, ACTN2, ZASP and titin Z1Z2 in the axial view. The Z-disc in cardiac muscle contains 4-6 ACTN2 repeats [5], and here 6 repeats are shown. The ACTN2 molecules can also be organised into doublets of the homodimer, spaced 6 nm apart ([13] not shown here). **B:** Arrangement of actin, ACTN2, ZASP and titin Z1Z2 in the transverse view (basketweave lattice). The lattice constants for titin Z1Z2 and parallel actin filaments are stable across the Z-disc, while the sets of anti-parallel actin filaments may move relative to one another. The rotation of the square lattice between different locations on the Z-disc is also shown [5].

A component describing repeated localisations of the same molecule was found in the best model for Z1Z2 and ZASP organisation, but not in the best ACTN2 model. The overall fit at the shortest distances is also worse for ACTN2. The most likely explanation for these two observations is the increased likelihood of nearby pairs of ACTN2 molecules in the Z-disc structure, which add an extra, unmodelled distance that confounds the fit of a zero-distance component. This could be consistent with the presence of pairs of ACTN2 dimers, seen in EM density maps [13], which are approximately 6 nm apart.

In the transverse plane (Fig. 2D-F), the most likely model for the arrangement of all three proteins included a characteristic distance of 27–30 nm, with a variability (σ_broadening_) of 7–9 nm (**Table 3**, **Fig. 3B**). This distance also corresponds to the spacing of parallel actin filaments (*d*_p-fil_) in this plane and the ACTN2 binding sites on them in the basketweave lattice structure [5]. It is also in broad agreement with our previous estimate for the transverse distance of 24 nm measured in fixed cardiomyocytes using 3D dSTORM followed by PERPL analysis [22]. Interestingly, the Z-discs in the cardiomyocytes were more likely to have adopted the small square lattice form, as the cells were predominantly relaxed when fixed, and thus the lattice spacing would be expected to be about 10-20% smaller.

For ACTN2, the best model also included a short characteristic distance at 10 ± 8 nm between localisations in the transverse plane. The Adhiron binds to the calponin homology (CH) domains of ACTN2, which are located either side of the actin filament. Thus, the 10 nm distance could arise from by binding of Adhirons to either side of the actin filament (**Fig 3A**). However, we do not have an explanation for an additional characteristic distance of 51 ± 8 nm, a small component in the best ACTN2 model (Table 3). We did not find a distance matching the spacing between anti-parallel actin filaments in the model, and the background term is a large contribution to the total RPD (**Fig. 2**D). This suggests that the transverse separation between ACTN2 binding sites (*d*_p-fil_) is relatively consistent between binding sites on parallel filaments in a square lattice structure, but less consistent between binding sites on anti-parallel filaments. A long-range variability of the displacement between the anti-parallel filament lattices may account for this (**Fig 3B**).

For Z1Z2, the best model also included distances at 1/√2 times (19 ± 7 nm; mean ± σ_broadening_, fitted s.d.) and √2 times (38 ± 7 nm) the repeating value of 27 ± 7 nm between localisations in the transverse plane. These distances could arise from distances between anti-parallel actin filaments (1/√2 × lattice constant) and between second-nearest neighbour parallel filaments (√2 × lattice constant) in the small square lattice structure. The second most likely model for the transverse arrangement of Z1Z2, which was 3.5 times less likely to describe the true pattern, also contained the distances at 19 nm and 38 nm (between anti-parallel filaments) but lacked the distance at 27 nm (between parallel filaments) (**Fig. S4**).

We conclude that the transverse distances between Z1Z2 titin domains are on a square lattice structure with a lattice constant of *d*_p-fil_ / √2 (**Fig 3B**).

Finally, for ZASP, the best model also included a second peak at twice the main distance of 28 ± 9 nm in the transverse plane, although it made a very small contribution to the total model RPD and its centre was beyond the maximum distance fitted. This peak may simply account for a very slightly higher density than a simple flat background at longer distance values. We therefore conclude that ZASP is most consistently found separated by the distance between parallel actin filaments in the transverse plane, similar to the ACTN2-actin binding site.

## Discussion

Identifying the organisation of Z-disc proteins, using a wide range of approaches, has proven to be very difficult. To improve on this, we labelled cardiac myofibrils using Adhirons, specific for Z-disc proteins, which have the advantage that their small size enables them to penetrate the Z-disc better than antibodies, and the linkage error between the target and the dye is only a few nm [20]. We imaged the fluorescent Adhirons using iPALM, which has a 2-5x improvement in resolution in x,y and z compared to other single molecule localisation techniques [17]. We then analysed the iPALM data using PERPL [22], to successfully extract repeating distances that likely reflect the underlying organisation of 3 different Z-disc proteins in cardiac myofibrils. We found a repeating pattern of approximately 16 nm for ACTN2 molecules along the actin filament in x,y, which is in reasonable agreement with electron microscopy data, in which ACTN2 molecules can be clearly observed [5, 13].

Likewise, the main repeat distance in x,y was about 16-18 nm for ZASP and Z1Z2. In the transverse plane, we found a repeat distance of about 27-30 nm for all three proteins, equivalent to the distance between parallel actin filaments. However, only the localisation data for Z1Z2 additionally showed distances equivalent to that for anti-parallel actin filaments. Overall, this suggests that the Z1Z2 domains have the highest degree of organisation in the Z-disc. The variability, or relative inconsistency, that we have inferred in the distance between ACTN2 binding domains on anti-parallel filaments may be related to the discrepancy between actin symmetry and perfect square lattice angles [9].

Our new results for ACTN2 reveal more detail than our previous study using PERPL to analyse dSTORM data [22, 30], in which the resolution was lower than achievable using iPALM. In particular, the improved axial resolution from iPALM allows us to resolve the repeating organisation of ACTN2 in the transverse plane, with a repeat distance of 27–30 nm, as well as a 10 nm repeat we ascribe to on binding sites either side of a single actin filament (Fig. 3). These results are consistent with recent work using cryo-electron tomography (cryo-ET) to visualise the Z-disc in mouse skeletal muscle that revealed ACTN2 doublets spaced about 6 nm apart, and up to 3 pairs of ACTN2 molecules, spanning about 37 nm along the actin filament, with a spacing of approximately 18.5 nm between the doublets [13]. While we cannot resolve the 6 nm separation with iPALM, our new data would be consistent with the presence of these doublets in cardiac muscle. The overall organisation of ACTN2 in the Z-disc was also reported to be less well ordered by CryoET than in previous models [13], again consistent with our data. The overall agreement of the PERPL data for ACTN2 with EM data suggests we can also be confident about our findings for other Z-disc proteins.

The linear repeat of 16-18 nm for the titin Z1Z2 domains has not been reported before. Previous reports have either located the N-terminal titin Z1Z2 domain to the central region or towards the outside of the Z-disc [26, 31–33]. The two Z1Z2 domains, each comprised of an Ig-like fold (4nm in size), form a highly stable complex in the Z-disc composed of antiparallel dimers between which t-cap (telethonin) is sandwiched [34–36]. The Z1Z2 domains are followed by a serine-proline rich linker region (Zis-1) and then up to seven Z-repeats (45 residues, 12 nm long) that may run through the Z-disc [25, 26, 37]. The number of Z-repeats is lower in fast (narrow Z-discs) than in slow and cardiac muscles (wider Z-discs), regulated through differential gene splicing of the Z-repeats [26, 38]. This led to the idea that z-repeat number regulates the number of ACTN2 molecules and hence width of the Z-disc. However, the span of each Z-repeat (12 nm) is lower than the spacing between ACTN2 molecules [39], only Z-repeats 1,3,5 and 7 bind to ACTN2 (C-terminal region), and this binding is weak [40].

Our results are not consistent with Z1Z2 only being located at the edge of the Z-disc. The regular spacing that we observe is unlikely to be explained by off-target effects, as the Z1Z2 Adhiron is specific, and does not bind to downstream Z-disc sequence. The dimensions of the axial and transverse patterns are consistent with a regular interaction of the N-terminal region of titin with ACTN2 throughout the Z-disc. It is possible that an interaction of Z2-Zis-1 with ACTN2 may contribute to this [37]. Importantly, the arrangement of Z1Z2 is more highly ordered than that of ACTN2, both in the axial and transverse directions, which suggests other interactions likely contribute and that the titin N-terminal region is a key organiser of the Z-disc.

The linear repeat of 16-18 nm that we also observed for ZASP is consistent with its binding to ACTN2. ZASP comprises an N-terminal PDZ domain followed by a downstream intrinsically disordered region that contains several LIM (Lin11, Isl-1, Mec-3) domains and a ZASP-like motif (ZM). The PDZ and ZM domains bind to the C-terminal region (150 residues) of ACTN2 [41]. Differential splicing results in three cardiac isoforms of ZASP, of which only two contain the 3 C-terminal LIM domains. The shorter isoform without LIM domains is only found in mature cardiac muscle and may block growth of the Z-disc [42–44]. The Adhiron to ZASP recognises the PDZ domain, present in all 3 ZASP isoforms in adult cardiac muscle. Our data suggests that there are fewer ZASP molecules than ACTN2 molecules in the Z-disc and that ZASP does not appear to be restricted to any specific region of the Z-disc.

Z-discs contain over 50 different proteins [1] in a narrow structure. Determining the location and organisation of each of these is a highly challenging task. Here our combined use of Adhirons, iPALM data and the use of PERPL for analysis has allowed us to obtain quantitative, protein-specific spatial analysis down to 10 nm, despite the low density of iPALM localisations, particularly for ZASP. PERPL is well-suited to such a task, where low density of localisations precludes the effective use of particle averaging approaches from fluorescence localisations [45, 46] and where distinguishing specific proteins by EM is not possible. Future work using single molecule localisation approaches combined with PERPL will help us to address the challenge of uncovering protein organisation in this complex structure.

## Materials and methods

### Adhirons – labelling and specificity

Three different Adhirons were used, raised against the calponin homology (CH) domains of ɑ-actinin-2 (ACTN2), the Z1Z2 repeats of titin, and the protein ZASP. We have briefly reported on these Adhirons previously [23]. Once isolated, the coding sequence is subcloned into a bacterial expression vector containing a unique C-terminal cysteine. The Adhirons were expressed in BL21 STAR^TM^ (DE3) *E. coli* (Life Technologies, Invitrogen) and affinity purified using Ni-NTA resin (Thermo Scientific) as previously described [23, 47]. Briefly, a single colony was used to inoculate a 2 ml overnight culture of LB media supplemented with 100 µg/mL carbenicillin. Then, 50 ml LB media plus antibiotic was inoculated with 1 ml of the overnight culture and grown at 37 °C and 230 rpm to an OD_600_ between 0.6–0.8. Protein production was induced by addition of IPTG 0.1 mM and incubated for a further 20–22 hr, at 25°C and 150 rpm before harvesting. His-tagged proteins were lysed in 1 ml Adhiron lysis buffer (50 mM NaH_2_PO_4_, 300 mM NaCl, 30 mM Imidazole, 10% Glycerol, 1% Triton X-100, pH 7.4) supplemented with 0.1 mg/ml lysozyme, Halt^TM^ protease inhibitor cocktail and 10 U/ml benzonase nuclease (Millipore, Burlington, MA). The lysate was then incubated with 200 µl of washed NiNTA His-Pur slurry (ThermoFisher Scientific) for 2 hr, washed with Adhiron wash buffer (50 mM NaH_2_PO_4_, 500 mM NaCl, 20 mM Imidazole, pH 7.4) and eluted in Adhiron elution buffer (50 mM NaH_2_PO_4_, 500 mM NaCl, 300 mM Imidazole, 20% glycerol, pH 7.4).

For iPALM, the Adhirons were labelled with AlexaFluor647, via a maleimide linkage to a unique cysteine residue at the C-terminus, and then further purified to remove extraneous dye as previously described [47]. Briefly, 150 µl of 40 µM Adhiron solutions were incubated with agitation at room temperature with 150 µl tris (2-carboxyethyl)phosphine (TCEP) immobilised resin (ThermoFisher Scientific) for 1 h. The solution was centrifuged (1 min at 1500 x *g*) and 130 µl of the supernatant transferred to a fresh tube containing 6 µl of 2 mM dye-maleimide (AlexaFluor-647-C2 maleimide, A20347 Invitrogen) and incubated at room temperature for 2 h. The reaction was quenched with 1.3 µl β-mercaptoethanol for 15 min at room temperature.

Excess label was removed using Zeba dye and biotin removal columns (Thermo Scientific) as per the manufacturer’s instructions, followed by dialysis into PBS.

### Target Protein Production

The calponin homology (CH) domains of ɑ-actinin-2 (ACTN2) were expressed and purified as previously described [48], the Z1Z2 repeats of titin (residues 1–200) [23], the end of Z2 repeat to the beginning of the Zr5 domain of titin (residues 201–598) and the protein ZASP PDZ (residues 1-89)[23] were subcloned in pGEX-6P-1 between the BamHI and EcoRI sites. Target proteins were expressed in *Escherichia coli* BL21 Rosetta 2 (Novagen) and purified using GST-tag affinity chromatography. Briefly, a single colony was used to inoculate a 7 ml overnight culture of 2YT media supplemented with 100 µg/mL carbenicillin and 34 µg/mL chloramphenicol. Then, 400 ml LB (ACTN2) or 500 ml TB media plus antibiotics was inoculated with 5 ml of the overnight culture and grown at 37 °C and 230 rpm to an OD600 between 0.6–0.8. Protein production was induced by addition of IPTG 0.5 mM and incubated for a further 3 h at 37 °C and 230 rpm or 2 h at 25 °C and 150 rpm before the temperature was reduced to 25°C for a further 16–18 h. Cells were harvested at 4000 x *g* at 4 °C for 20 mins. Harvested pellets were lysed in GST lysis buffer (50 mM Tris-HCl, 500mM NaCl, 1 mM DTT, 1 mM EDTA, 1% Triton X-100, pH 7.4) supplemented with 1x HALT_TM_ protease inhibitor cocktail, 1 mg/ml lysozyme and 3U/mL benzonase followed by sonication (6 cycles of 10 s on/off pulses), centrifuged (20,000 x *g*, 30 min). Lysates were then incubated with 1.25 mL washed Glutathione Resin (Amintra) for 1hr. Resin was washed five times with GST Wash buffer (50 mM Tris-HCl, 500 mM NaCl, 1 mM DTT, 1 mM EDTA, pH8.0) and proteins eluted with GST Elution buffer (50 mM Tris-HCl, 500 mM NaCl, 1 mM DTT, 1 mM EDTA, 20 mM reduced glutathione, pH 7.4).

### Immunoprecipitation and immunoblotting

Immunoprecipitations used His-Tag Dynabeads (ThermoFisher). Dynabeads were incubated with 50 µg Adhirons in 1x casein blocking buffer (SigmaAldrich) in wash buffer (100 mM sodium-phosphate, pH 8.0, 600 mM NaCl, 0.02% Tween-20) for 10 min and rinsed with wash buffer. Beads were then incubated with target proteins at a 1:1 molar ratio for 1 h at room temperature. Following three washes, proteins were eluted by incubation in His elution buffer (300 mM Imidazole, 50 mM sodium phosphate, pH 8.0, 300 mM NaCl, 0.01% Tween-20) for 10 min. Immunoprecipitants were heated in 4x SDS-sample buffer (200 mM Tris-HCl, 8% SDS, 20% glycerol, 10% mercaptoethanol, 0.1% (w/v) bromophenol blue, pH 7) and run on a 15% SDS-PAGE gel at 120V before transfer to nitrocellulose membrane using the BioRad Transblot Turbo. Membranes were then blocked in 5% milk in tris-buffered saline – 0.1% Tween 20 (TBS-T) incubation with rabbit anti-GST-HRP (1:10,000; Abcam, ab3416) or rabbit anti-6xHisTag-HRP (1:10,000; Abcam, ab1187) for 1 h at room temperature. Membranes were rinsed three times with TBS-T before development using Immunoblot Forte Western HRP (Millipore), according to the manufacturer’s instructions. Blots were imaged using an Amersham_TM_ Imager 600 (GE Healthcare, Chicago, IL).

### Sample preparation

Frozen pig hearts were obtained from Pel-Freez Biologicals. The heart was allowed to thaw, small pieces (approximately 200-300 mg each) were excised from the left ventricular muscle wall using a scalpel and used to generate myofibrils using a similar approach to those used to generate mouse heart myofibrils [49]. Briefly, ∼250 mg of muscle was placed into 1 ml of K60 buffer + BDM in an eppendorf and rinsed 2-3 times until the buffer was mostly clear.

K60 buffer contains 1 x Protease inhibitor cocktail (Sigma P8340). The muscle was then transferred into 1 ml of K60 buffer in a 5 ml Eppendorf tube on ice and homogenised using a tissue tearor, at max speed (∼21K RPM) for ∼30s, repeated twice until no visible intact tissue remained. The mixture was then transferred to a 1.7 ml Eppendorf tube, and centrifuged at 4°C, 1000 g for 10 minutes. The supernatant was removed, and the pellet resuspended in K60 buffer containing 1% Triton X-100, then incubated on ice for 30 minutes. The mixture was vortexed twice during this incubation. The mixture was then centrifuged at 4°C, 1000 g for 5 minutes, the supernatant removed and the pellet resuspended in 1 ml K60 buffer. This mixture was then centrifuged at 4°C, 1000 g for 5 minutes, the supernatant removed, and the pellet resuspended in K60 buffer supplemented with 0.1% BSA, 10mM DTT and 1 mM EGTA. If not used immediately, myofibrils were stored for up to a week in the final K60 buffer containing 50% glycerol at -20°C. We thank David Warshaw’s group for sharing this detailed protocol with us [49].

To stain myofibrils, ∼50 µl of resuspended myofibrils in K60 buffer (containing 0.1% BSA) were added to a 25 mm diameter round coverslip (CS-25R17, #1.5 thickness; Warner Instruments) embedded with gold nanorod fiducial markers (A12-40-600-CTAB; Nanopartz) and allowed to attach for ∼ 2 minutes. If using myofibrils stored in glycerol, ∼50 µl was removed and placed in a 1.7 ml Eppendorf, 450 µl of PBS was added, and the mixture was centrifuged at 2000 rpm for 5 minutes at 4°C. The supernatant was removed, and the pellet resuspended in 50 µl of PBS containing 0.1% BSA and then added to the coverslip. The coverslip was washed gently once with phosphate buffered saline (PBS), before fixing the myofibrils using 4% paraformaldehyde in PBS for 5 minutes. The coverslip was washed once with PBS, excess liquid removed, and PBS containing 2% BSA was added to the coverslip (blocking step) for 10 minutes. Excess solution was removed, and the fluorescent Adhiron added, diluted 1/10 to 1/20 depending on the concentration of the Adhiron, in PBS containing 0.2% Triton X 100. The coverslip was incubated at room temperature with the Adhiron for ∼20-30 minutes at room temperature, or overnight at 4°C. The coverslip was then washed twice with PBS containing 0.2% Triton X 100 and briefly stored in PBS before mounting in the imaging chamber.

17mg/mL Catalase (C100-50MG, Sigma-Aldrich) and 70mg/mL Glucose Oxidase (G-2133-50KU, Sigma-Aldrich) solutions were prepared in Buffer A (10mM Tris pH 8.0 + 50mM NaCl). Both were mixed in a 1:4 volume ratio to prepare GLOX solution. This was centrifuged and only the supernatant was used. 1M cysteamine (30070-50G, Sigma-Aldrich) solution was prepared in 0.25N HCl. The STORM buffer was prepared by combining GLOX solution, 1M cysteamine, and Buffer B (50mM Tris pH 8.0 + 10mM NaCl + 10% Glucose) in 1:10:90 volume ratio. The sample coverslip was covered with STORM-buffer. An 18 mm diameter round coverslip (CS-18R17, #1.5 thickness; Warner Instruments) was placed centrally on top of the sample and the boundary was sealed with Valap (https://cshprotocols.cshlp.org/content/2015/2/pdb.rec082917) to create a sandwich mounting.

### Sample imaging

iPALM imaging was performed similarly to as described in previous work [14, 17, 50–52]. The Alexa Fluor 647-labelled samples were excited using 647 nm laser (Opto Engine LLC) excitation at ca. 2–3 kW/cm^2^ intensity in TIRF conditions. The gold fiducials embedded in the coverslip were used for calibrating interferometry. 100,000 images were collected through dual Nikon Apo TIRF 60x/1.49NA objective lenses coupled to a 647 nm long-pass filter (LP02-647RU, Semrock), and acquired via three EMCCD cameras (DU987E, Andor) at 30 ms exposure. The field-of-view (FOV) varied from 30-40 µm.

### iPALM data preprocessing

As described in [14, 17, 50–52], the iPALM data was processed/localized using PeakSelector software (Janelia Research Campus). The gold fiducials embedded in the coverslip were used for drift correction. We used the grouped xyz position output as final localisation coordinates. z is depth through the sample along the optical axis.

### Z-disc segmentation

Z-discs were segmented from iPALM reconstructions as reported previously [29]. Briefly, the point cloud data from each FOV was rendered as a 3D histogram with 50-nm bin size in x, y and z. The Z-discs in each image were then segmented using Ilastik’s pixel classification and object classification workflow, implemented through the graphical user interface (see Supplementary Videos 1-8) [28]. The underlying localisations (point-cloud data) for each Z-disc were then extracted from these segmented histograms. Z-disc dimensions and volume were then calculated respectively using principal component analysis (PCA) and the convex hull of the localisations. Sarcomere length was measured using the line selection tool in Fiji.

### PERPL modelling

The organisation of each protein in the Z-disc in both the axial direction (cell axis, y) and in the transverse plane (xz) was modelled using PERPL [22] (**Fig. S2**). Briefly, PERPL calculates the relative position distribution (RPD) for the proteins, fits RPD models generated from hypothetical models of organisation in real space and gives relative likelihoods for these models. Each model RPD is constructed by summing terms for: the background from randomly distributed localisations, N_peaks_ number of peaks for the characteristic distances of type, d_type_, between the localisations; and the independent RPD resulting from repeated localisation of the same dye molecule (repeats).

The point-cloud for each Z-disc was pre-processed in preparation for PERPL analysis. First, the line of each Z-disc in its xy view was aligned along x using 2D PCA, leaving z unchanged. Next, density-based spatial clustering of applications with noise (DBSCAN) was applied to each Z-disc to remove outlier localisations far from the main body of the Z-disc, implemented in visualisation software Open3D [36, 53]. The two parameters for DBSCAN, epsilon (ε) and minimum points (minPts), were separately optimised for each Z-disc between ε: 50 – 150 nm and minPts: 3 – 7 by visually assessing the clustering result.

To generate the experimental RPD, the data was filtered by removing localisations with x, y or z precision above a threshold, σ_loc-max_. The Euclidean distances between the resulting localisations were calculated up to 150 nm in the direction or plane being modelled. No limit was placed on the distances between localisations in the orthogonal directions, as this significantly reduced the number of distances available for modelling. From this set of distances, the final RPD was calculated as a kernel density estimate using Churchman’s distributions for distances between localisations in 1D or 2D [54], up to a maximum distance, L_fit_. To minimise the impact of edge effects in the transverse direction, L_fit_ was set well below the extent of the data in xz (**Tables 1,** Table **S2**).

Background terms were a function of the relative distance (d) between localisations, including linear_1D_ (Ad + B, A ≤ 0, B ≥ 0), linear_flat_ (Ad + B, A ≤ 0, B ≥ 0 for 0 ≤ d < -B/A; 0 for d ≥ -B/A), linear_2D_ (Ad + B, A ≥ 0, B ≥ 0), linear_2D,int=0_ (Ad, A ≥ 0), quad_2D_ (Ad + Bd^2^, A ≥ 0, B ≤ 0), cubic_2D_ (Ad + Bd^2^ + Cd^3^, A ≥ 0, B ≤ 0, C ≥ 0), None (no background term) or constant background level. A, B and C were optimised during fitting. 2D background options were extensions to PERPL based on the theoretical distribution of distances for randomly sampled points within a finite rectangle [55].

Peak type in the model RPD, d_type_, was “int”, where peaks were at integer multiples of a single characteristic distance, *a*; “sq”, where peaks were at *a*, *a*√2, 2*a*; or “ind”, where characteristic distances were independent of one another. These peaks have a width defined by a Gaussian-like broadening term, σ_broadening_, which is optimised for each model (rather than per-peak) during model fitting, to reflect experimental noise and biological variability [54] [22] . The characteristic distance values were optimised during model fitting. The term for repeat localisations of the same molecule (zero-distance) has a separate σ_broadening_.

Model RPDs were fit to each experimental RPD by adjusting the model parameters to minimise the mean-squared error (MSE), having specified initial guesses and bounds for the parameter values. In the transverse plane, the count in the experimental and model RPD was divided by the distance, to avoid the fit being dominated by a parameter for a linearly increasing count as expected for a random 2D distribution (ignoring edge effects) [22, 30].

We extended PERPL to allow the model RPDs to be procedurally generated. This allowed us to sweep over multiple pre-processing parameters and models to explore variability and inform final model selection in each case (**Table S1, Supplementary Spreadsheet**). Pre-processing filters were strengthened until remaining data was insufficient to generate a usable experimental RPD. For each protein and 3D model component (axial or transverse), we selected the lowest σ_loc-max_ (most precise localisations) at which models appeared to fit well and the longest L_fit_ for final model selection (**Tables 1, S2, Supplementary Spreadsheet**).

For each experimental RPD, we selected the model with the lowest corrected Akaike information criterion (AIC_c_) as the most likely, as in previous work [22]. AIC_c_ measures differences in information loss between different models for the same data, and therefore the relative likelihood that the models describe the true distribution from which the data is sampled. It penalises overfitting:

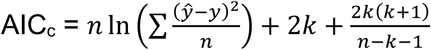

for models fitted by minimising MSE, where *n* is the number of datapoints, *k* is the number of free parameters, *y* is the experimental RPD and *ŷ* is the model RPD [56]. Models that had a relative likelihood ≥ 0.01 compared with the most likely model (ΔAIC_c_ ≤ 9.21) were also identified [56]. Models with a fitted parameter value at its permitted bounds, negative background (not physically real) or large uncertainty on the parameter value (parameter uncertainty > parameter value) were rejected. AIC_c_ may not be used to compare models fitted to different data, so between different localisation filtering settings.

## Acknowledgements

iPALM experiments were performed in collaboration with the Advanced Imaging Center (AIC) at Janelia Research Campus, a facility supported the Howard Hughes Medical Institute. We would like to thank Harry Takagi, Pauline Bennett and Dave Warshaw for help and advice with the protocols used to generate and stain myofibrils.

## Data and resource availability

The iPALM localisation data is available at https://doi.org/10.5281/zenodo.19661651.

Analysis used the software at https://github.com/oubino/z_disk/releases/tag/v0.0.2 with the version of PERPL at https://pypi.org/project/perpl/. The scripts used for PERPL modelling are also included in https://github.com/AlistairCurd/PERPL-Python3/releases/tag/zdisc-2.0.

## Supplementary Material

**Figure S1.**
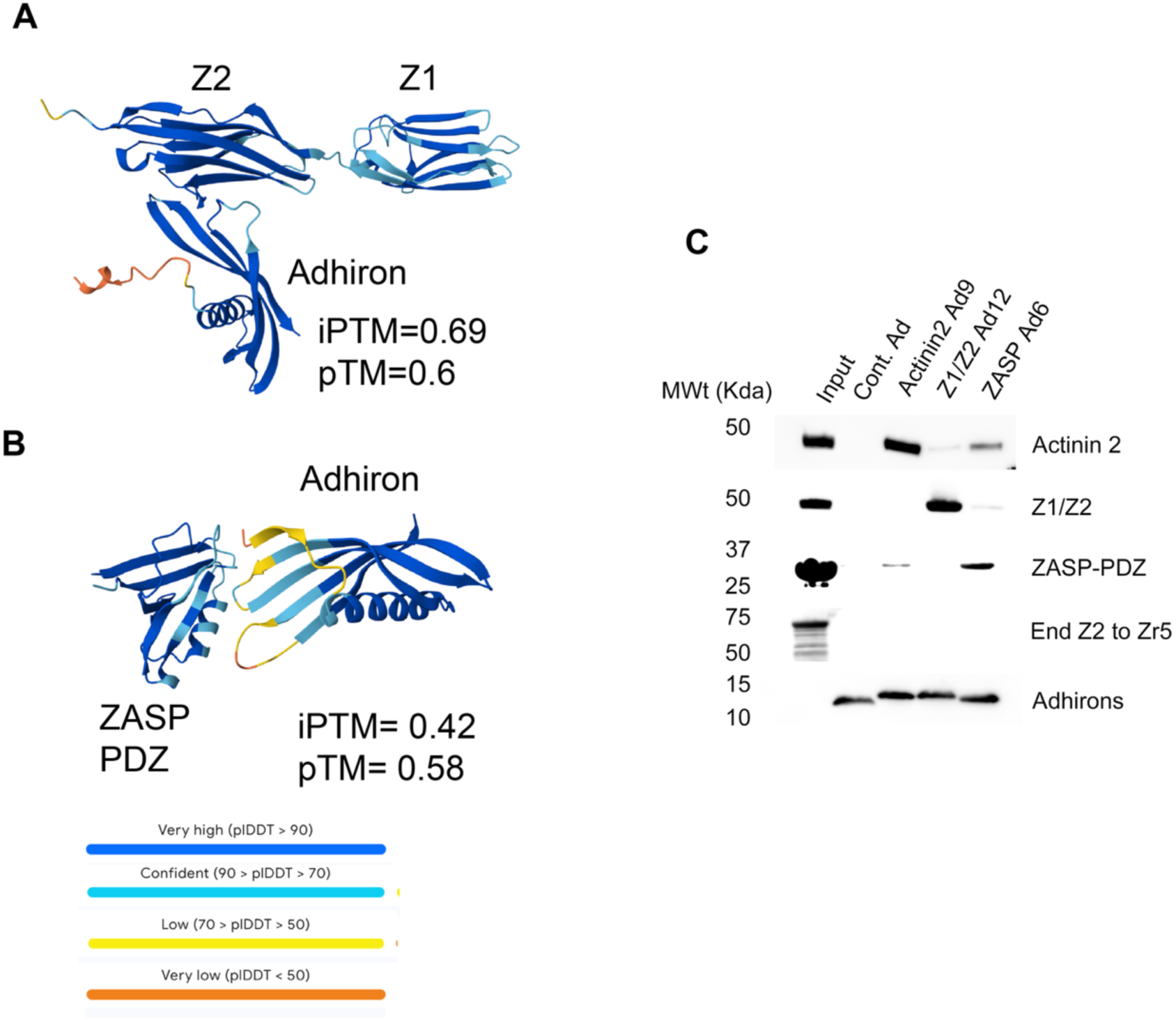
Z-disc Adhirons are specific for their targets. Alphafold modelling shows the potential interaction of the Adhirons for Z1Z2 (**A**) and ZASP (**B**) with their target proteins.The specificity of the Z-disc Adhirons for their target proteins was confirmed by immunoprecipitations using the Adhiron His-Tag, His-Tag Dynabeads and GST-tagged target proteins, followed by immunoblotting for GST and His Tags. All Adhirons bound their targets with minimal cross-reactivity and no Adhirons bound to the End Z2 to Zr5 Titin protein. Ad – Adhiron, Cont. – control (an Adhiron with AAAA and AAE in the variable loops). Representative blots shown. N=3.

**Figure S2.**
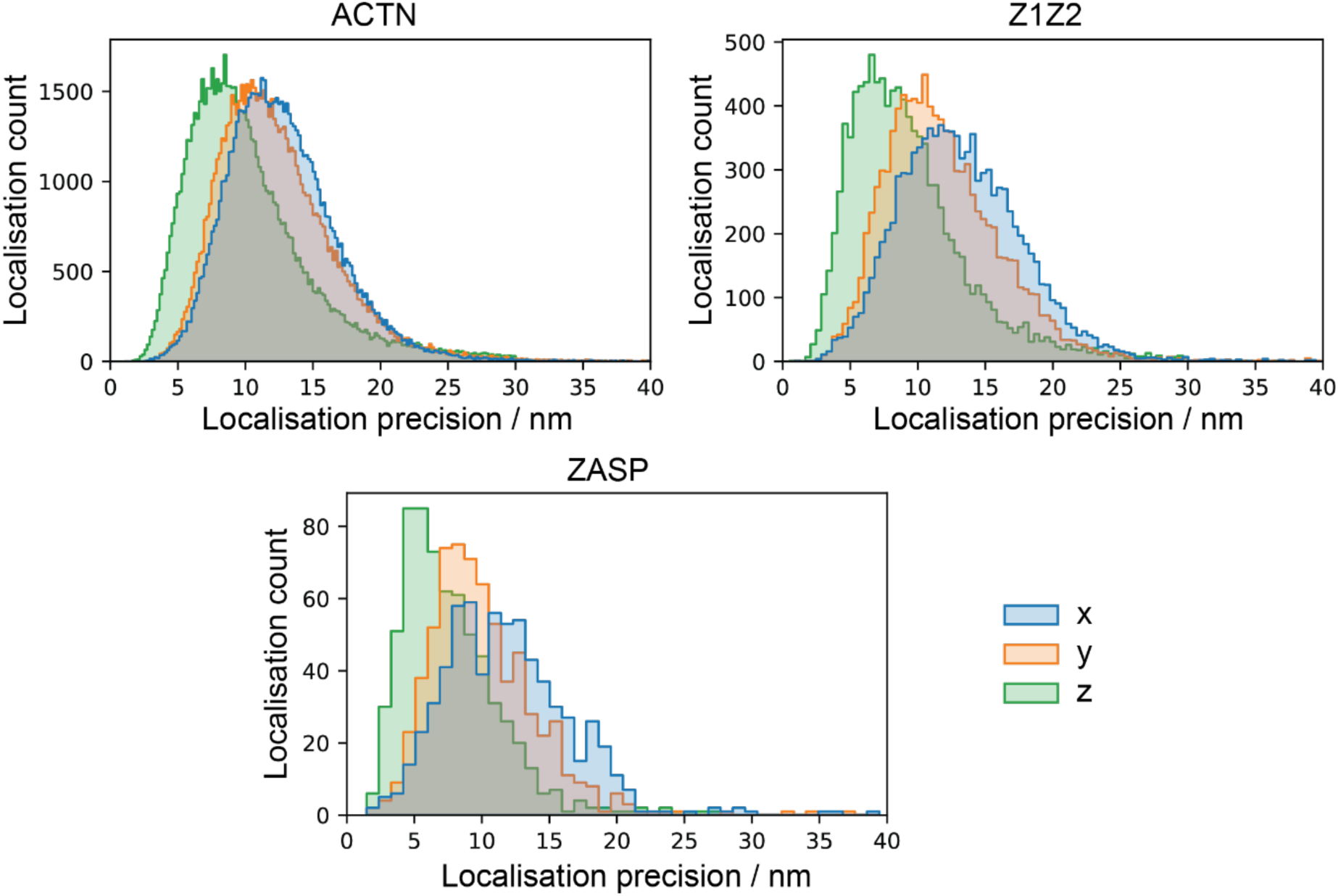
Histograms for the localisation precision in x, y and z for each protein before filtering by localisation precision or number of localisations per Z-disc.

**Figure S3.**
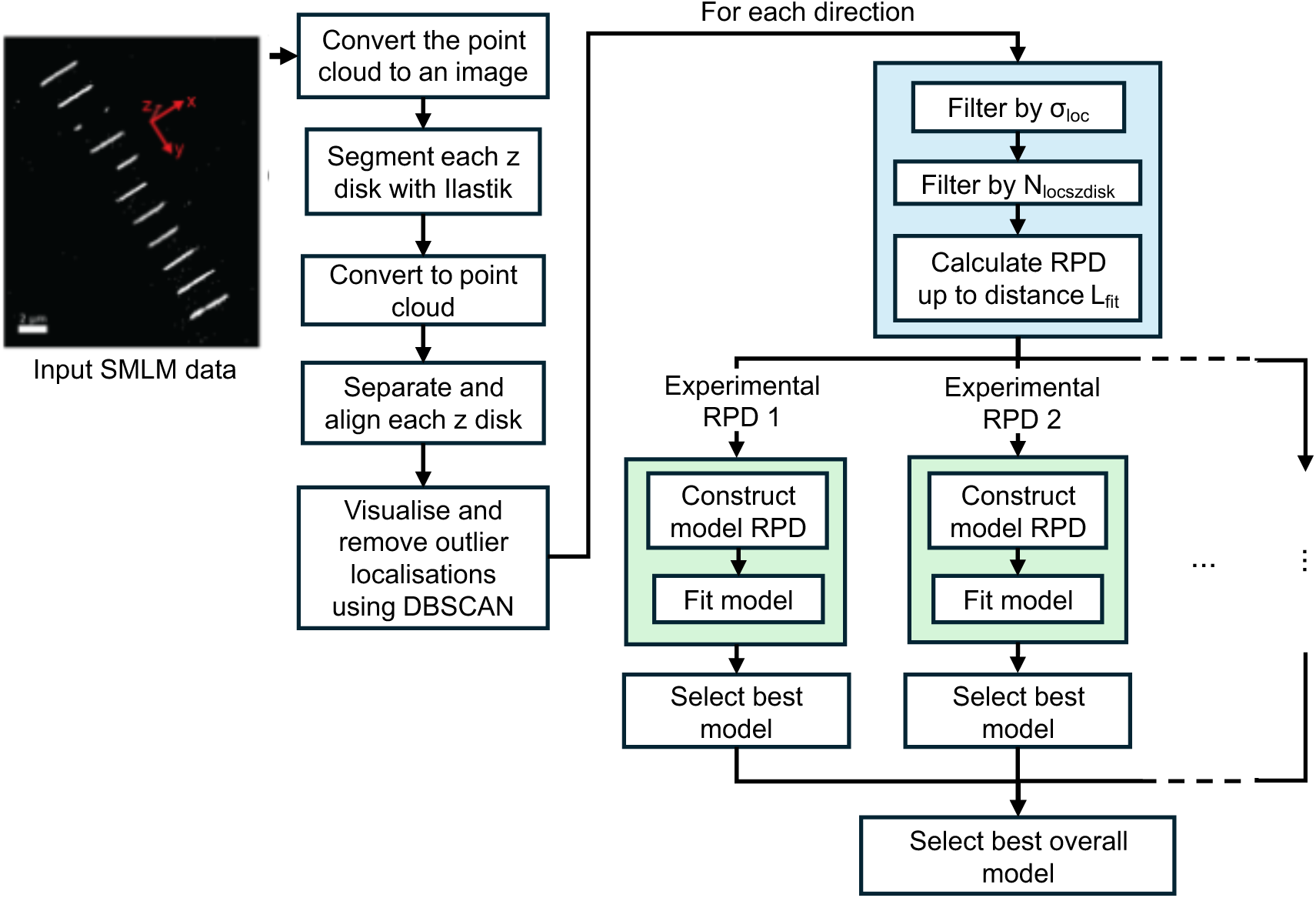
Analysis workflow applied to the SMLM data for each protein. Light blue box: sweep over range of values for σ_loc_, N_locszdisc_ and L_fit_ to generate multiple experimental RPDs. Light green box: sweep over range of possible models.

**Figure S4.**
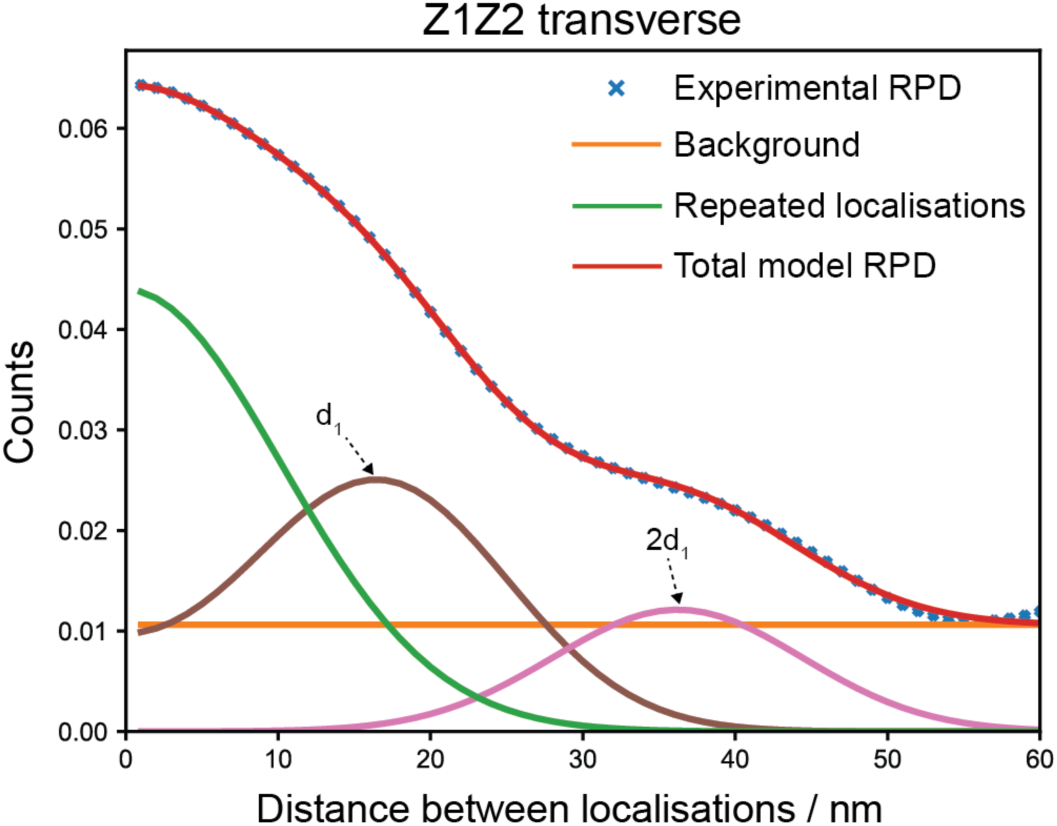
Next most likely model for Z1Z2 in the transverse plane. Characteristic distance d_1_: 18.57 ± 0.07 nm with σ_broadening_: 8.09 ± 0.08 nm. Experimental and model RPD parameters are σ_loc_: 7 nm, L_fit_: 60 nm, N_locszdisc_: 1, background: Linear_2D,int=0_, N_peaks_: 2, d_type_: int, repeats: True, N_distances_: 134.

**Table S1.** Hyperparameter sweep. Parameter values used in the exhaustive sweep over possible experimental RPDs and models for each protein and direction combination. σ_loc_: maximum estimated localisation precision filter. N_locszdisc_: minimum number of localisations in a Z-disc for inclusion of the Z-disc. L_fit_: Distance over which the model was fitted. N_peaks_: number of characteristic distances included in the model RPD. d_type_: Relationship between characteristic distances in the model (**Methods**). Repeats: Inclusion of a term for repeated localisations of a single molecule.

|  |  | Experimental RPD |  |  | Model |  |  |  |
| --- | --- | --- | --- | --- | --- | --- | --- | --- |
| Protein | Direction | $\sigma_{loc}$ (nm) | $N_{locszdisc}$ | $L_{fit}$ (nm) | Background | $N_{peaks}$ | $d_{type}$ | Repeats |
| ACTN2 | Axial | 5, 6, 7 | 1, 5, 10 | 105, 115, 125 | None, Flat, Linear<sub>1D</sub>, Linear<sub>flat</sub> | 0, 1, 4, 5, 6 | int | False, True |
| ZASP | Axial | 9, 10 | 1 | 110, 115, 120 | None, Flat, Linear<sub>1D</sub>, Linear<sub>flat</sub> | 0, 1, 4, 5, 6 | int | False, True |
| Z1Z2 | Axial | 7, 8, 9 | 1, 5, 10 | 100, 110, 120 | None, Flat, Linear<sub>1D</sub>, Linear<sub>flat</sub> | 0, 1, 4, 5, 6 | int | False, True |
| ACTN2 | Transverse | 5, 6, 7 | 1, 5, 10 | 52, 57, 62 | None, Flat, Linear<sub>2D</sub>, Linear<sub>2D,int=0</sub>, Quad<sub>2D</sub>, Cubic<sub>2D</sub> | 0, 2, 3 | int, sq, ind | False, True |
| ZASP | Transverse | 9, 10 | 1 | 42, 47, 52 | None, Flat, Linear<sub>2D</sub>, Linear<sub>2D,int=0</sub>, Quad<sub>2D</sub>, Cubic<sub>2D</sub> | 0, 2, 3 | int, sq, ind | False, True |
| Z1Z2 | Transverse | 7, 8, 9 | 1, 5, 10 | 50, 55, 60 | None, Flat, Linear<sub>2D</sub>, Linear<sub>2D,int=0</sub>, Quad<sub>2D</sub>, Cubic<sub>2D</sub> | 0, 2, 3 | int, sq, ind | False, True |

**Table S2.** Pre-processing parameters for final experimental RPD. Filters used to obtain the experimental RPD for final model selection for each protein and direction.

| **Protein** | **Direction** | **$\sigma_{\text{loc}}$ / nm** | **$N_{\text{locszdisc}}$** | **$L_{\text{fit}}$ / nm** | **$N_{\text{distances}}$** |
| --- | --- | --- | --- | --- | --- |
| ACTN2 | Axial | 6 | 1 | 125 | 457 |
| Z1Z2 | Axial | 8 | 1 | 120 | 406 |
| ZASP | Axial | 9 | 1 | 120 | 69 |
| ACTN2 | Transverse | 6 | 1 | 62 | 462 |
| Z1Z2 | Transverse | 7 | 1 | 60 | 134 |
| ZASP | Transverse | 9 | 1 | 52 | 69 |

**Table S3.**
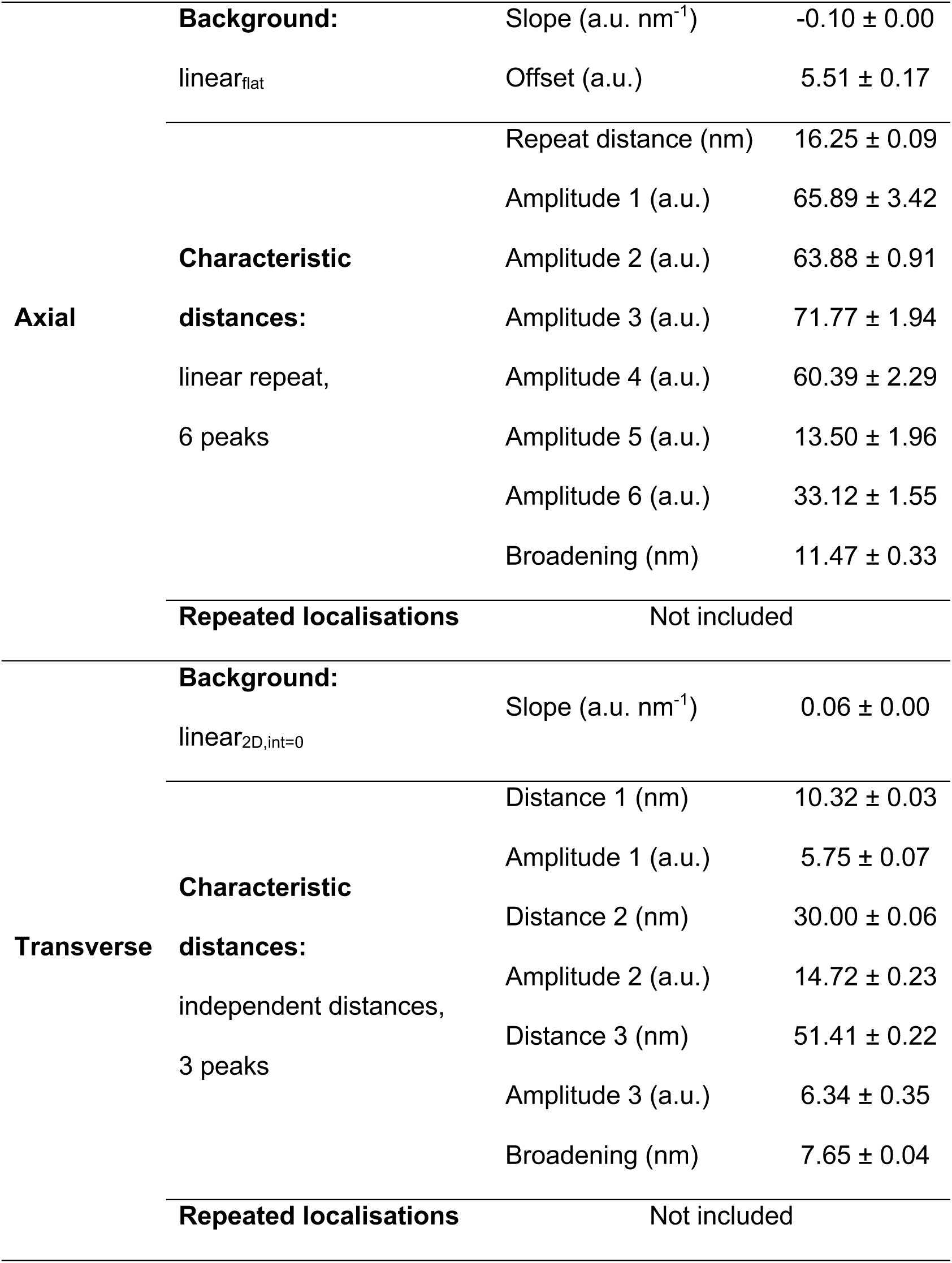
Most likely models and parameter values for ACTN2 arrangement.

**Table S4.**
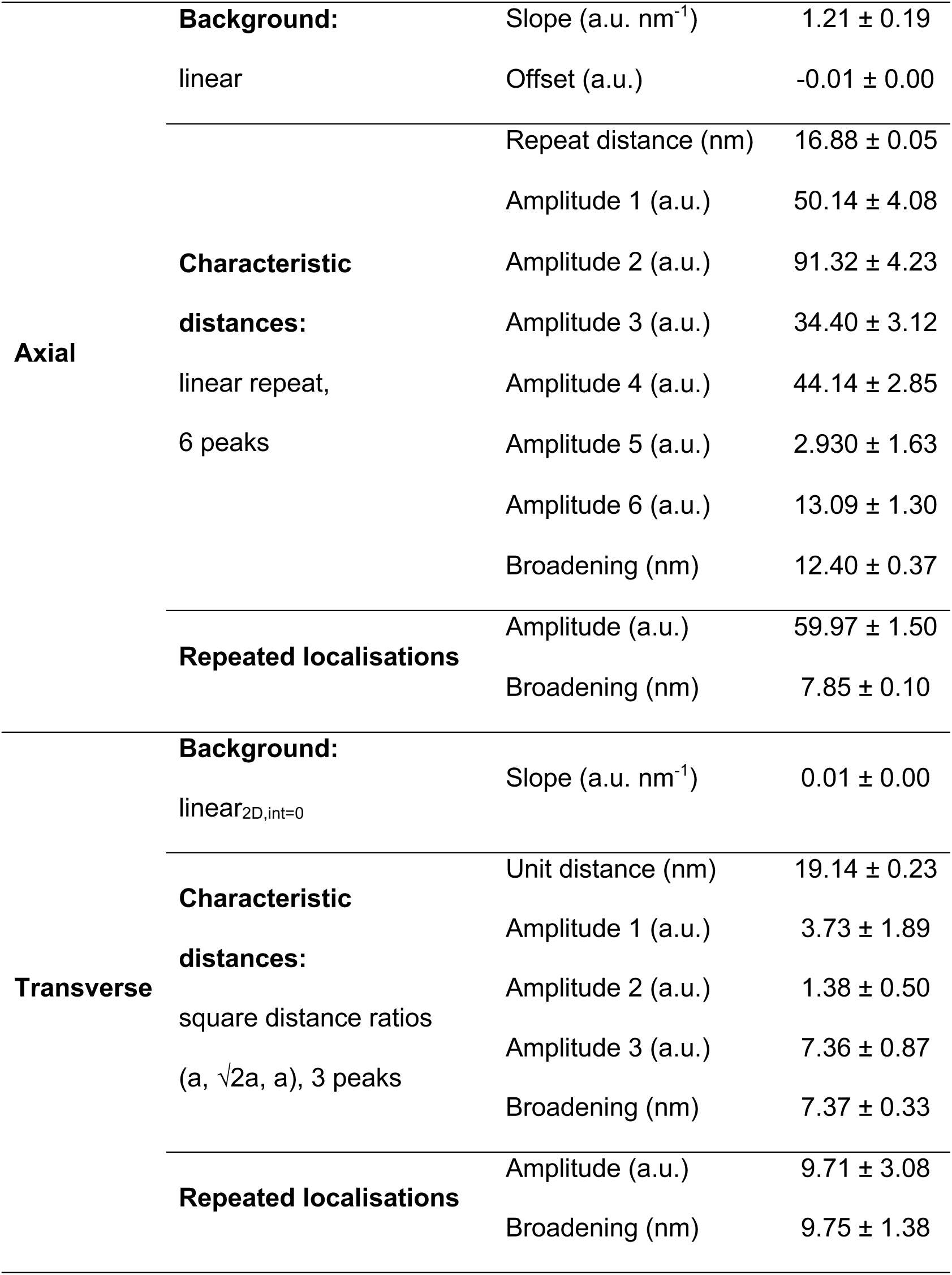
Most likely models and parameter values for Z1Z2 arrangement.

**Table S5.**
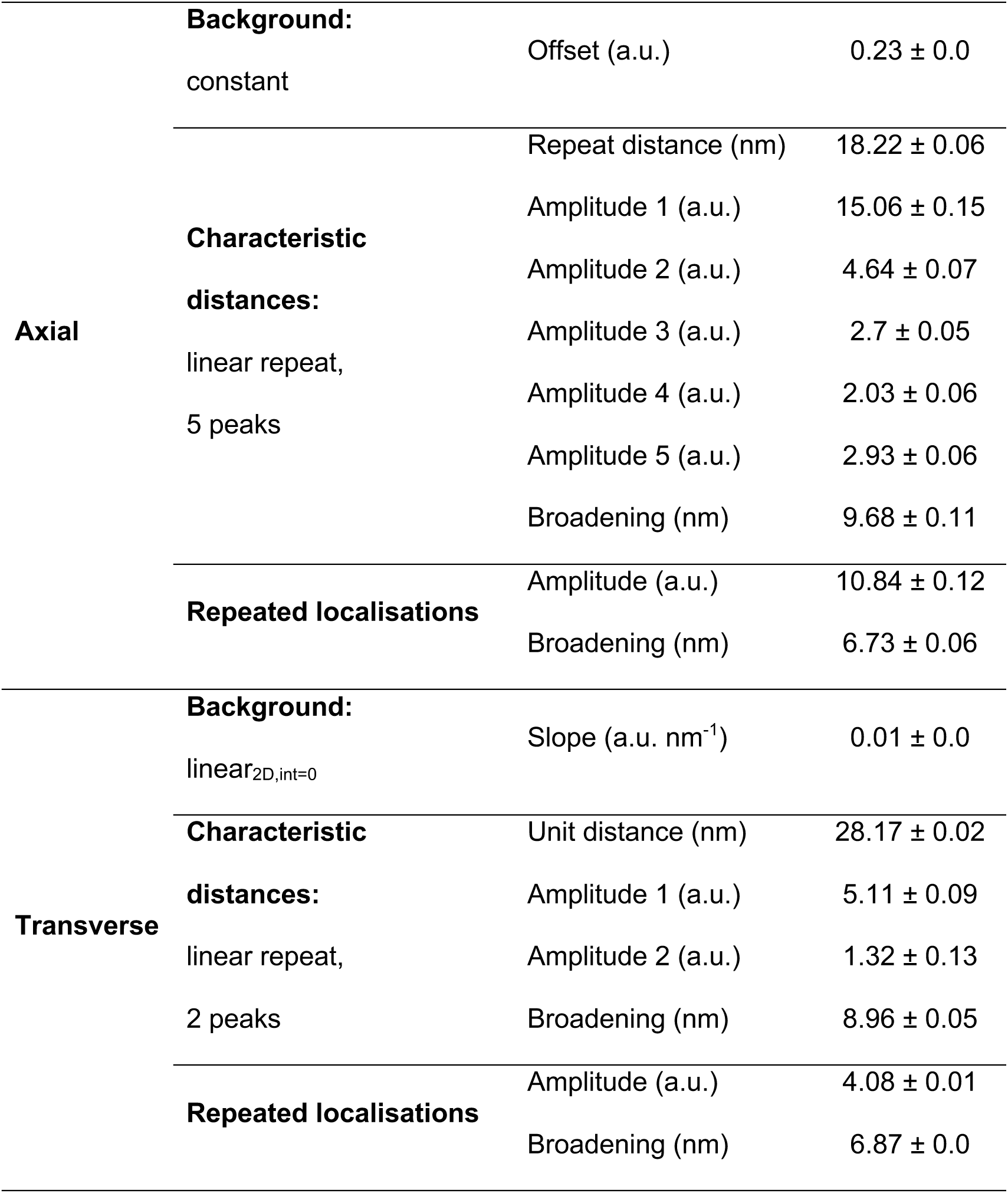
Most likely models and parameter values for ZASP arrangement.

## Supplementary Spreadsheet

(see separate Excel file)

This Spreadsheet includes all of the models that successfully arrived at a fit for all of the pre-processing options.

## References

1. Reimann L, Wiese H, Leber Y, Schwable AN, Fricke AL, Rohland A, et al. Myofibrillar Z-discs Are a Protein Phosphorylation Hot Spot with Protein Kinase C (PKCalpha) Modulating Protein Dynamics. Mol Cell Proteomics. 2017;16(3):346–67. Epub 20161227. doi: 10.1074/mcp.M116.065425. PubMed PMID: 28028127; PubMed Central PMCID: PMCPMC5340999.

2. Frank D, Frey N. Cardiac Z-disc signaling network. J Biol Chem. 2011;286(12):9897–904. Epub 20110121. doi: 10.1074/jbc.R110.174268. PubMed PMID: 21257757; PubMed Central PMCID: PMCPMC3060542.

3. Wadmore K, Azad AJ, Gehmlich K. The Role of Z-disc Proteins in Myopathy and Cardiomyopathy. Int J Mol Sci. 2021;22(6). Epub 20210317. doi: 10.3390/ijms22063058. PubMed PMID: 33802723; PubMed Central PMCID: PMCPMC8002584.

4. Schock F, Gonzalez-Morales N. The insect perspective on Z-disc structure and biology. J Cell Sci. 2022;135(20). Epub 20221013. doi: 10.1242/jcs.260179. PubMed PMID: 36226637.

5. Luther PK. The vertebrate muscle Z-disc: sarcomere anchor for structure and signalling. J Muscle Res Cell Motil. 2009;30(5-6):171–85. Epub 20091015. doi: 10.1007/s10974-009-9189-6. PubMed PMID: 19830582; PubMed Central PMCID: PMCPMC2799012.

6. Goldstein MA, Michael LH, Schroeter JP, Sass RL. Structural states in the Z band of skeletal muscle correlate with states of active and passive tension. J Gen Physiol. 1988;92(1):113–9. doi: 10.1085/jgp.92.1.113. PubMed PMID: 3171533; PubMed Central PMCID: PMCPMC2228892.

7. Perz-Edwards RJ, Reedy MK. Electron microscopy and x-ray diffraction evidence for two Z-band structural states. Biophys J. 2011;101(3):709–17. doi: 10.1016/j.bpj.2011.06.024. PubMed PMID: 21806939; PubMed Central PMCID: PMCPMC3145273.

8. Yamaguchi M, Izumimoto M, Robson RM, Stromer MH. Fine structure of wide and narrow vertebrate muscle Z-lines. A proposed model and computer simulation of Z-line architecture. J Mol Biol. 1985;184(4):621–43. doi: 10.1016/0022-2836(85)90308-0. PubMed PMID: 4046026.

9. Oda T, Yanagisawa H. Cryo-electron tomography of cardiac myofibrils reveals a 3D lattice spring within the Z-discs. Commun Biol. 2020;3(1):585. Epub 20201016. doi: 10.1038/s42003-020-01321-5. PubMed PMID: 33067529; PubMed Central PMCID: PMCPMC7567829.

10. Edwards RJ, Goldstein MA, Schroeter JP, Sass RL. The Z-band lattice in skeletal muscle in rigor. J Ultrastruct Mol Struct Res. 1989;102(1):59–65. doi: 10.1016/0889-1605(89)90033-5. PubMed PMID: 2621377.

11. Burgoyne T, Morris EP, Luther PK. Three-Dimensional Structure of Vertebrate Muscle Z-Band: The Small-Square Lattice Z-Band in Rat Cardiac Muscle. J Mol Biol. 2015;427(22):3527–37. Epub 20150908. doi: 10.1016/j.jmb.2015.08.018. PubMed PMID:

12. 26362007; PubMed Central PMCID: PMCPMC4641244.

12. Rusu M, Hu Z, Taylor KA, Trinick J. Structure of isolated Z-disks from honeybee flight muscle. J Muscle Res Cell Motil. 2017;38(2):241–50. Epub 20170721. doi: 10.1007/s10974-017-9477-5. PubMed PMID: 28733815; PubMed Central PMCID: PMCPMC5660141.

13. Wang Z, Grange M, Wagner T, Kho AL, Gautel M, Raunser S. The molecular basis for sarcomere organization in vertebrate skeletal muscle. Cell. 2021;184(8):2135–50 e13. Epub 20210324. doi: 10.1016/j.cell.2021.02.047. PubMed PMID: 33765442; PubMed Central PMCID: PMCPMC8054911.

14. Kanchanawong P, Shtengel G, Pasapera AM, Ramko EB, Davidson MW, Hess HF, et al. Nanoscale architecture of integrin-based cell adhesions. Nature. 2010;468(7323):580–4. doi: 10.1038/nature09621. PubMed PMID: 21107430; PubMed Central PMCID: PMCPMC3046339.

15. Mund M, van der Beek JA, Deschamps J, Dmitrieff S, Hoess P, Monster JL, et al. Systematic Nanoscale Analysis of Endocytosis Links Efficient Vesicle Formation to Patterned Actin Nucleation. Cell. 2018;174(4):884–96 e17. Epub 20180726. doi: 10.1016/j.cell.2018.06.032. PubMed PMID: 30057119; PubMed Central PMCID: PMCPMC6086932.

16. Liu S, Hoess P, Ries J. Super-Resolution Microscopy for Structural Cell Biology. Annu Rev Biophys. 2022;51:301–26. Epub 20220204. doi: 10.1146/annurev-biophys-102521-112912. PubMed PMID: 35119945.

17. Shtengel G, Galbraith JA, Galbraith CG, Lippincott-Schwartz J, Gillette JM, Manley S, et al. Interferometric fluorescent super-resolution microscopy resolves 3D cellular ultrastructure. Proc Natl Acad Sci U S A. 2009;106(9):3125–30. Epub 20090206. doi: 10.1073/pnas.0813131106. PubMed PMID: 19202073; PubMed Central PMCID: PMCPMC2637278.

18. Fruh SM, Matti U, Spycher PR, Rubini M, Lickert S, Schlichthaerle T, et al. Site-Specifically-Labeled Antibodies for Super-Resolution Microscopy Reveal In Situ Linkage Errors. ACS Nano. 2021;15(7):12161–70. Epub 20210629. doi: 10.1021/acsnano.1c03677. PubMed PMID: 34184536; PubMed Central PMCID: PMCPMC8320235.

19. Carrington G, Tomlinson D, Peckham M. Exploiting nanobodies and Affimers for superresolution imaging in light microscopy. Mol Biol Cell. 2019;30(22):2737–40. Epub 2019/10/15. doi: 10.1091/mbc.E18-11-0694. PubMed PMID: 31609674; PubMed Central PMCID: PMCPMC6789155.

20. Cordell P, Carrington G, Curd A, Parker F, Tomlinson D, Peckham M. Affimers and nanobodies as molecular probes and their applications in imaging. J Cell Sci. 2022;135(14). Epub 20220718. doi: 10.1242/jcs.259168. PubMed PMID: 35848463; PubMed Central PMCID: PMCPMC9450889.

21. Tiede C, Tang AA, Deacon SE, Mandal U, Nettleship JE, Owen RL, et al. Adhiron: a stable and versatile peptide display scaffold for molecular recognition applications. Protein Eng Des Sel. 2014;27(5):145–55. doi: 10.1093/protein/gzu007. PubMed PMID: 24668773; PubMed Central PMCID: PMC4000234.

22. Curd AP, Leng J, Hughes RE, Cleasby AJ, Rogers B, Trinh CH, et al. Nanoscale Pattern Extraction from Relative Positions of Sparse 3D Localizations. Nano Lett. 2021;21(3):1213–20. Epub 20201130. doi: 10.1021/acs.nanolett.0c03332. PubMed PMID: 33253583; PubMed Central PMCID: PMCPMC7883386.

23. Parker F, Tang AAS, Rogers B, Carrington G, Dos Remedios C, Li A, et al. Affimers targeting proteins in the cardiomyocyte Z-disc: Novel tools that improve imaging of heart tissue. Front Cardiovasc Med. 2023;10:1094563. Epub 20230214. doi: 10.3389/fcvm.2023.1094563. PubMed PMID: 36865889; PubMed Central PMCID: PMCPMC9971620.

24. Tskhovrebova L, Trinick J. Titin: properties and family relationships. Nat Rev Mol Cell Biol. 2003;4(9):679–89. doi: 10.1038/nrm1198. PubMed PMID: 14506471.

25. Gregorio CC, Trombitas K, Centner T, Kolmerer B, Stier G, Kunke K, et al. The NH2 terminus of titin spans the Z-disc: its interaction with a novel 19-kD ligand (T-cap) is required for sarcomeric integrity. J Cell Biol. 1998;143(4):1013–27. doi: 10.1083/jcb.143.4.1013. PubMed PMID: 9817758; PubMed Central PMCID: PMCPMC2132961.

26. Gautel M, Goulding D, Bullard B, Weber K, Furst DO. The central Z-disk region of titin is assembled from a novel repeat in variable copy numbers. J Cell Sci. 1996;109 ( Pt 11):2747–54. doi: 10.1242/jcs.109.11.2747. PubMed PMID: 8937992.

27. Faulkner G, Pallavicini A, Formentin E, Comelli A, Ievolella C, Trevisan S, et al. ZASP: a new Z-band alternatively spliced PDZ-motif protein. J Cell Biol. 1999;146(2):465–75. doi: 10.1083/jcb.146.2.465. PubMed PMID: 10427098; PubMed Central PMCID: PMCPMC3206570.

28. Berg S, Kutra D, Kroeger T, Straehle CN, Kausler BX, Haubold C, et al. ilastik: interactive machine learning for (bio)image analysis. Nat Methods. 2019;16(12):1226–32. Epub 20190930. doi: 10.1038/s41592-019-0582-9. PubMed PMID: 31570887.

29. Umney O, Leng J, Canettieri G, Galdo NAR, Slaney H, Quirke P, et al. Annotation and automated segmentation of single-molecule localisation microscopy data. J Microsc. 2024;296(3):214–26. Epub 20240802. doi: 10.1111/jmi.13349. PubMed PMID: 39092628.

30. Curd A, Cleasby A, Baird M, Peckham M. Modelling 3D supramolecular structure from sparse single-molecule localisation microscopy data. J Microsc. 2024;296(2):115–20. Epub 20231025. doi: 10.1111/jmi.13236. PubMed PMID: 37877157.

31. Atkinson RA, Joseph C, Kelly G, Muskett FW, Frenkiel TA, Nietlispach D, et al. Ca2+-independent binding of an EF-hand domain to a novel motif in the alpha-actinin-titin complex. Nat Struct Biol. 2001;8(10):853–7. doi: 10.1038/nsb1001-853. PubMed PMID: 11573089.

32. Yajima H, Ohtsuka H, Kawamura Y, Kume H, Murayama T, Abe H, et al. A 11.5-kb 5’-terminal cDNA sequence of chicken breast muscle connectin/titin reveals its Z line binding region. Biochem Biophys Res Commun. 1996;223(1):160–4. doi: 10.1006/bbrc.1996.0862. PubMed PMID: 8660363.

33. Furst DO, Osborn M, Nave R, Weber K. The organization of titin filaments in the half-sarcomere revealed by monoclonal antibodies in immunoelectron microscopy: a map of ten nonrepetitive epitopes starting at the Z line extends close to the M line. J Cell Biol. 1988;106(5):1563–72. doi: 10.1083/jcb.106.5.1563. PubMed PMID: 2453516; PubMed Central PMCID: PMCPMC2115059.

34. Bertz M, Wilmanns M, Rief M. The titin-telethonin complex is a directed, superstable molecular bond in the muscle Z-disk. Proc Natl Acad Sci U S A. 2009;106(32):13307–133310. Epub 20090721. doi: 10.1073/pnas.0902312106. PubMed PMID: 19622741; PubMed Central PMCID: PMCPMC2726412.

35. Lee EH, Gao M, Pinotsis N, Wilmanns M, Schulten K. Mechanical strength of the titin Z1Z2-telethonin complex. Structure. 2006;14(3):497–509. doi: 10.1016/j.str.2005.12.005. PubMed PMID: 16531234.

36. Zhou Q-Y, Park J, Koltun V. Open3D: A Modern Library for 3D Data Processing. arxivorg/abs/180109847. 2018.

37. Labeit S, Lahmers S, Burkart C, Fong C, McNabb M, Witt S, et al. Expression of distinct classes of titin isoforms in striated and smooth muscles by alternative splicing, and their conserved interaction with filamins. J Mol Biol. 2006;362(4):664–81. Epub 20060801. doi: 10.1016/j.jmb.2006.07.077. PubMed PMID: 16949617.

38. Sorimachi H, Freiburg A, Kolmerer B, Ishiura S, Stier G, Gregorio CC, et al. Tissue-specific expression and alpha-actinin binding properties of the Z-disc titin: implications for the nature of vertebrate Z-discs. J Mol Biol. 1997;270(5):688–95. doi: 10.1006/jmbi.1997.1145. PubMed PMID: 9245597.

39. Luther PK, Squire JM. Muscle Z-band ultrastructure: titin Z-repeats and Z-band periodicities do not match. J Mol Biol. 2002;319(5):1157–64. doi: 10.1016/S0022-2836(02)00372-8. PubMed PMID: 12079354.

40. Grison M, Merkel U, Kostan J, Djinovic-Carugo K, Rief M. alpha-Actinin/titin interaction: A dynamic and mechanically stable cluster of bonds in the muscle Z-disk. Proc Natl Acad Sci U S A. 2017;114(5):1015–20. Epub 20170117. doi: 10.1073/pnas.1612681114. PubMed PMID: 28096424; PubMed Central PMCID: PMCPMC5293040.

41. Au Y, Atkinson RA, Guerrini R, Kelly G, Joseph C, Martin SR, et al. Solution structure of ZASP PDZ domain; implications for sarcomere ultrastructure and enigma family redundancy. Structure. 2004;12(4):611–22. doi: 10.1016/j.str.2004.02.019. PubMed PMID: 15062084.

42. Sheikh F, Bang ML, Lange S, Chen J. "Z"eroing in on the role of Cypher in striated muscle function, signaling, and human disease. Trends Cardiovasc Med. 2007;17(8):258–62. doi: 10.1016/j.tcm.2007.09.002. PubMed PMID: 18021935; PubMed Central PMCID: PMCPMC2134983.

43. Huang C, Zhou Q, Liang P, Hollander MS, Sheikh F, Li X, et al. Characterization and in vivo functional analysis of splice variants of cypher. J Biol Chem. 2003;278(9):7360–5. Epub 20021223. doi: 10.1074/jbc.M211875200. PubMed PMID: 12499364.

44. Fisher LAB, Schock F. The unexpected versatility of ALP/Enigma family proteins. Front Cell Dev Biol. 2022;10:963608. Epub 20221201. doi: 10.3389/fcell.2022.963608. PubMed PMID: 36531944; PubMed Central PMCID: PMCPMC9751615.

45. Wu Y-L, Hoess P, Tschanz A, Matti U, Mund M, Ries J. Maximum-likelihood model fitting for quantitative analysis of SMLM data. Nature Methods. 2023;20(1):139–48. doi: 10.1038/s41592-022-01676-z.

46. Heydarian H, Joosten M, Przybylski A, Schueder F, Jungmann R, Werkhoven Bv, et al. 3D particle averaging and detection of macromolecular symmetry in localization microscopy. Nature Communications. 2021;12(1):2847. doi: 10.1038/s41467-021-22006-5.

47. Tiede C, Bedford R, Heseltine SJ, Smith G, Wijetunga I, Ross R, et al. Affimer proteins are versatile and renewable affinity reagents. Elife. 2017;6. Epub 20170627. doi: 10.7554/eLife.24903. PubMed PMID: 28654419; PubMed Central PMCID: PMCPMC5487212.

48. Haywood NJ, Wolny M, Rogers B, Trinh CH, Shuping Y, Edwards TA, et al. Hypertrophic cardiomyopathy mutations in the calponin-homology domain of ACTN2 affect actin binding and cardiomyocyte Z-disc incorporation. Biochem J. 2016;473(16):2485–93. doi: 10.1042/BCJ20160421. PubMed PMID: 27287556; PubMed Central PMCID: PMC4980809.

49. Nelson S, Beck-Previs S, Sadayappan S, Tong C, Warshaw DM. Myosin-binding protein C stabilizes, but is not the sole determinant of SRX myosin in cardiac muscle. J Gen Physiol. 2023;155(4). Epub 20230123. doi: 10.1085/jgp.202213276. PubMed PMID: 36688870; PubMed Central PMCID: PMCPMC9884578.

50. Case LB, Baird MA, Shtengel G, Campbell SL, Hess HF, Davidson MW, et al. Molecular mechanism of vinculin activation and nanoscale spatial organization in focal adhesions. Nat Cell Biol. 2015;17(7):880–92. Epub 20150608. doi: 10.1038/ncb3180. PubMed PMID: 26053221; PubMed Central PMCID: PMCPMC4490039.

51. Stubb A, Guzman C, Narva E, Aaron J, Chew TL, Saari M, et al. Superresolution architecture of cornerstone focal adhesions in human pluripotent stem cells. Nat Commun. 2019;10(1):4756. Epub 20191018. doi: 10.1038/s41467-019-12611-w. PubMed PMID: 31628312; PubMed Central PMCID: PMCPMC6802214.

52. Kumari R, Ven K, Chastney M, Kokate SB, Peranen J, Aaron J, et al. Focal adhesions contain three specialized actin nanoscale layers. Nat Commun. 2024;15(1):2547. Epub 20240321. doi: 10.1038/s41467-024-46868-7. PubMed PMID: 38514695; PubMed Central PMCID: PMCPMC10957975.

53. Ester M, Kriegel H-P, Sander J, Xu X. A density-based algorithm for discovering clusters in large spatial databases with noise. Proceedings of the Second International Conference on Knowledge Discovery and Data Mining; Portland, Oregon: AAAI Press; 1996. p. 226–31.

54. Churchman LS, Flyvbjerg H, Spudich JA. A non-Gaussian distribution quantifies distances measured with fluorescence localization techniques. Biophys J. 2006;90(2):668–71. Epub 20051028. doi: 10.1529/biophysj.105.065599. PubMed PMID: 16258038; PubMed Central PMCID: PMCPMC1367071.

55. Mathai AM, Moschopoulos P, Pederzoli G. Random points associated with rectangles. Rendiconti del Circolo Matematico di Palermo. 1999;48(1):163–90. doi: 10.1007/BF02844387.

56. Burnham KP, Anderson DR. Model Selection and Multimodel Inference: A Practical Information-Theoretic Approach: Springer New York; 2003.

